# CXCR4 allows T cell acute lymphoblastic leukemia to escape from JAK1/2 and BCL2 inhibition through CNS infiltration

**DOI:** 10.1101/734913

**Authors:** Kirsti L. Walker, Sean P. Rinella, Nicholas J. Hess, David P. Turicek, Sabrina A. Kabakov, Fen Zhu, Myriam N. Bouchlaka, Sydney L. Olson, Monica M. Cho, Aicha E. Quamine, Arika S. Feils, Tara B. Gavcovich, Lixin Rui, Christian M. Capitini

**Affiliations:** Department of Pediatrics, University of Wisconsin School of Medicine and Public Health, 1111 Highland Avenue, Madison, WI 53705 USA; Carbone Cancer Center, University of Wisconsin School of Medicine and Public Health, 1111 Highland Avenue, Madison, WI 53705 USA; Department of Medicine, University of Wisconsin School of Medicine and Public Health, 1111 Highland Avenue, Madison, WI 53705 USA

**Keywords:** T cell acute lymphoblastic leukemia, ruxolitniib, venetoclax, plerixafor, JAK/STAT, BCL2

## Abstract

Targeting the JAK/STAT and BCL2 pathways in patients with relapsed/refractory T cell acute lymphoblastic leukemia (T-ALL) may provide an alternative approach to achieve clinical remissions. Ruxolitinib and venetoclax show a dose-dependent effect on T-ALL individually, but combination treatment reduces survival and proliferation of T-ALL in vitro. Using a xenograft model, the combination treatment fails to improve survival, with death from hind limb paralysis. Despite on-target inhibition by the drugs, histopathology demonstrates increased leukemic infiltration into the central nervous system (CNS) as compared to liver or bone marrow. Liquid chromatography-tandem mass spectroscopy shows that ruxolitinib and venetoclax insufficiently cross into the CNS. The addition of the CXCR4 inhibitor plerixafor with ruxolitinib and venetoclax reduces clinical scores and enhances survival. While combination therapy with ruxolitinib and venetoclax shows promise for treating T-ALL, additional inhibition of the CXCR4-CXCL12 axis may be needed to maximize the possibility of complete remission.

## Introduction

T cell acute lymphoblastic leukemia (T-ALL) accounts for 25% of adult and 15% of pediatric acute lymphoblastic leukemia (ALL) cases. T-ALL occurs when mutated thymocytes that are destined for cell death survive and acquire additional mutations that promote full malignant transformation[1]. Cure rates, respectively, reach 60% in adults and over 80% in pediatric patients when treated with combination chemotherapy protocols[2-4]. However, resistance to first-line therapy is seen in 25% of children and more than 50% of adults, and the cohorts that do respond to current ALL chemotherapy protocols often suffer from both acute and chronic toxic side-effects despite treatment success.[5] Therefore, there is a need for the development of novel, non-chemotherapeutic treatments for this type of blood cancer[6,7].

In order to improve prognosis and mitigate side-effects seen with current chemotherapy treatments, it is crucial to evaluate the underlying molecular pathways that drive T-ALL and contribute to chemotherapy resistance. Over the last decade, the Janus Activating Kinase/ Signal Transducer and Activator of Transcription (JAK/STAT) pathway has been to shown to play a critical pathogenic role in several hematologic malignancies including T-ALL[8,9]. Activation of JAK tyrosine kinases leads to the subsequent phosphorylation and activation of STAT transcription factors[10-12]. Additionally, many T-ALL express elevated levels of the pro-survival protein BCL2[13-15]^6^. These pathways allow for increased T-ALL proliferation through the JAK/STAT pathway and increased T-ALL survival through augmented BCL2 expression.

Ruxolitinib is an FDA approved small molecule inhibitor that targets JAK1 and JAK2. Ruxolitinib prevents the tyrosine phosphorylation of STAT1/STAT3/STAT5, which are downstream of cytokine receptors that drive T-ALL proliferation (e.g. IL-7), and should function to inhibit the proliferative properties seen in T-ALL. Unfortunately for most patients, chronic therapy with ruxolitinib results in reactivation of JAK/STAT signaling and prevents T-ALL remission[16]. In these cases, BCL2 may also be hijacked and upregulated to promote survival of these cancer cells.

Venetoclax is a highly potent and specific BCL2 mimetic that acts as an artificial sensitizer, binding to the BH3 groove of the BCL2 protein without targeting platelets. This causes the downregulation of BCL2 and eventual cell death of T-ALL[17-19]. The combination of ruxolitinib and venetoclax is emerging as a viable treatment option for relapsed disease. These drugs have been utilized previously to treat IL-7 receptor alpha (IL7R*α*)-mutated T-ALL, which prolongs survival in a preclinical model but ultimately all animals succumb to leukemia[20]. Because the central nervous system (CNS) is a common site of T-ALL involvement, [2,21] [2,21] [2,21] [2,21] [2,21] [2,21] [2,21] [2,21] one potential limitation of inhibiting these pathways *in vivo* is that it is unknown if these drugs can cross the blood-brain barrier (BBB)^2,21^. Also, the mechanism by which T-ALL would escape the combination of ruxolitinib and venetoclax is entirely unknown.

The CXCR4-CXCL12 axis has been implicated as a potential pathway that drives T-ALL invasion into the CNS[22-25]. This study will demonstrate that ruxolitinib and venetoclax are efficacious *in vitro* to treat T-ALL, but are not effective *in vivo* by showing for the first time their inability to effectively cross the blood-brain-barrier (BBB) and treat T-ALL in the CNS. By deleting the CXCR4 gene from T-ALL or blocking its activity with plerixafor, prolonged survival was observed *in vivo* with decreased total and neurologic clinical scores. Thus, in the setting of CXCR4 inhibition, T-ALL CNS infiltration can be blocked and allow for the systemic clearance of T-ALL by ruxolitinib and venetoclax.

## Methods

### Tumor cell lines

Jurkat (TIB-152) clone E6-1 and Loucy (CRL-2629) (ATCC Manassas, VA) were used as human T-ALL cell lines. Jurkat-GFP cell line was generated by transfecting 2×10^5^ Jurkat cells using the Lonza 4D-Nucleofector X Unit (Basel, Switzerland). The Nucleofector was used to deliver 1μg of px330 plasmid encoding for Cas9 protein and a gRNA for the AAVS1 locus, as well as 1μg of an AAVS1-CAGGS-EGFP plasmid to serve as the donor template. The Nucleofection was carried out in SE buffer and the pulse code for the transfection was CL-120. The cells were cultured in RPMI post nucleofection and sorted on a FACSAria for GFP^+^ cells. Cells were cultured in RPMI 1640 1x medium supplemented with 10% heat-inactivated fetal bovine serum (FBS) (Gemini bio-products, Sacramento, CA), L-Glutamine (2 mM), penicillin/streptomycin (100 μ/ml) at 37°C, 5% CO_2_ and 95% humidity. Excluding FBS, all other media ingredients were purchased from Corning, Corning, NY. All cell lines were mycoplasma tested and authenticated prior to use (IDEXX Bioresearch, Westbrook, ME).

### In vivo T-ALL xenograft model

NOD/SCID/*Il2rg*^*tmwjl*^/Szj (NSG) breeder mice were purchased from Jackson Labs (Bar Harbor, ME) and bred internally at the Biomedical Research Model Services Breeding Core at University of Wisconsin-Madison. Mice were housed in accordance with the Guide for the Care and Use of Laboratory Mice and experiments were performed under an IACUC approved animal protocol. Both male and female mice aged 8-16 weeks were randomized into control or treatment groups. On Day +0, 2×10^6^ Jurkat cells in 200 μl of 1x PBS were injected intravenously into NSG mice to generate a xenograft T-ALL mouse model. Using flow cytometric analysis, at least 1% anti-human CD45 cancer burden was documented in the peripheral blood (usually day 5-7) before initiating treatments. Mice were orally gavaged once daily for 14-days with vehicle(s), 100μl 30 mg/kg/day ruxolitinib, 100μl 35 mg/kg/day venetoclax, or combination. Mice were monitored for survival, % weight change, and clinical symptoms of disease (activity, hunch, and hind-limb paralysis) using a modified scoring system (Supplemental Table 1).

### Ruxolitinib and venetoclax treatments

Ruxolitinib (INCB018424) and venetoclax (ABT-199) were purchased in powdered form (Active Biochemicals, Kowloon Bay, Kowloon, Hong Kong, Cat # A-1134, A-1231). For *in vitro* use, ruxolitinib was resuspended in 100% DMSO at a concentration of 100 mg/ml and diluted to a working stock of 50 μM (final concentration of DMSO being 10%). For *in vivo* use, ruxolitinib in 100% DMSO was added to 5% N,N-dimethylacetamide (Cat # D137510-500ml, Sigma Aldrich, St. Louis, MO) + 0.5% methylcellulose (Cat # ME136, Spectrum Chemical, New Brunswick, NJ) in H2O. For *in vitro* use, venetoclax was resuspended in 100% warm DMSO at a concentration of 50 mg/ml and diluted to a working stock of 200 nM (final concentration of DMSO being 11%). For *in vivo* use, venetoclax was resuspended in 100% EtOH at a concentration of 100 mg/ml. Appropriate amount of venetoclax was resuspended in 10% -100% EtOH + 30% PEG 400 (Cat # 1008415, Rigaku, The Woodlands, TX) + 60% Phosal 50 (Lipoid, Ludwigshafen, Germany).

For other assays, please see Supplementary Methods.

### Statistical methods

Statistics were performed using GraphPad Prism version 7.0 for the Macintosh OS (GraphPad Software, San Diego, CA). Data were expressed as mean ± SEM. For analysis of three or more groups, a non-parametric ANOVA test was performed with the Bonferroni or Sidak’s multiple comparisons post-test. Analysis of differences between two normally distributed test groups was performed using a two-sided Mann Whitney test. A p-value less than 0.05 was considered statistically significant.

## Results

### JAK/STAT and BCL2 are dysregulated in T-ALL

To confirm that both STATs and BCL2 are dysregulated in T-ALL, we first performed flow cytometric analysis of two T-ALL cell lines, Jurkat and Loucy, and compared their expression of pSTAT1, pSTAT3 and BCL2 against two healthy controls. The T-ALL lines exhibited a complete dysregulation of pSTAT expression with decreased levels of pSTAT1 and increased levels of pSTAT3 compared to healthy controls (Figure 1A-B). We did not detect any expression of pSTAT5 in either T-ALL line or healthy control (data not shown). Additionally, BCL2 was upregulated in the T-ALL lines (Figure 1C). We next did a secondary analysis of RNA-Seq data from a Children’s Oncology Group study of T-ALL patients[26] and analyzed STAT1, STAT3 and BCL2 expression upon initial diagnosis or relapse. We observed that STAT expression remained constant even after relapse, while the expression of BCL2 decreased, suggesting that these pathways are important for T-ALL survival (Figure 1D-F).

**Figure 1.**
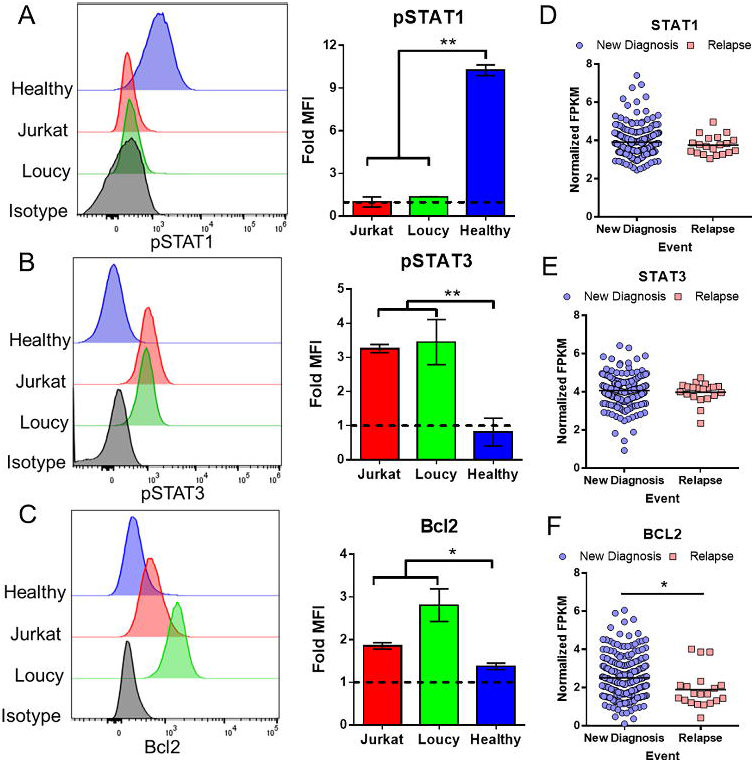
JAK/STAT and BCL2 pathways are dysregulated in T-ALL. (A-C) Histograms of T-ALL lines Jurkat and Loucy stained for pSTAT1 (A), pSTAT3 (B) or BCL2 (C) and compared to healthy controls and an isotype control antibody. Bar graphs represent the fold MFI (target MFI / isotype MFI) of two independent experiments (for cancer lines) and two healthy donors. (D-F) Normalized FPKM values of STAT1 (D), STAT3 (E) and BCL2 (F) at diagnosis and relapse from a Children’s Oncology Group T-ALL RNA-seq dataset[26].

### Single agent and combination inhibitor responses of ruxolitinib and venetoclax decrease the survival and proliferation of T-ALL cells *in vitro*

To investigate the relevance of the JAK/STAT and BCL2 pathways on T-ALL proliferation and cell survival, Jurkat (a mature T-ALL) and Loucy (an early precursor T-ALL) were assessed following treatment with either ruxolitinib or venetoclax.[15,27] Both of these cell lines were treated with a serial dose of ruxolitinib and venetoclax over the course of 72 h and evaluated using a trypan blue exclusion assay and MTT proliferation assay. Both ruxolitinib or venetoclax were able to decrease the survival and proliferation of both Jurkat and Loucy cell lines after 24, 48, and 72 h of treatment in a dose-dependent manner (Figure 2 and Supplemental Figure 1-2).

**Figure 2.**
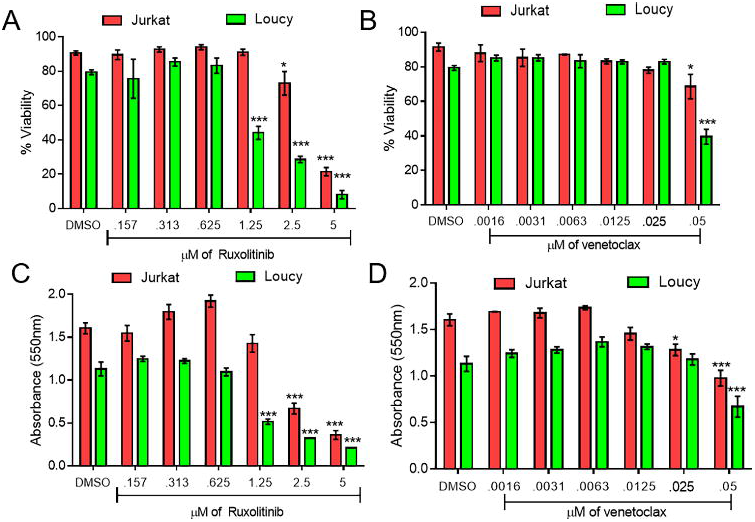
Potency of single-agent ruxolitinib and venetoclax against T-ALL lines. Trypan blue (A-B) and MTT proliferation data (C-D) of serially diluted ruxolitinib (A, C) and venetoclax (B, D) at 48 hours. Two independent experiments were performed with triplicate measurements. * p < 0.05; ** p < 0.01; *** p < 0.001.

After determining that ruxolitinib and venetoclax individually affect survival and proliferation of T-ALL, T-ALL cells were treated with ruxolitinib and venetoclax in combination and evaluated again for survival and proliferation. This combination dose was then tested for survival and proliferation in comparison to single-dose and DMSO controls in both Jurkat and Loucy cell lines (Figure 3A-B, Supplemental Figure 3). We also assessed both Annexin V and 7-AAD via flow cytometry and found increased Annexin V and 7-AAD in both the Jurkat and Loucy cell lines (Figure 3C). Lastly, the Jurkat cell line also showed a reduction in survival for up to 5 days in culture utilizing Incucyte live-cell imaging technology (p< 0.0001 compared to no treatment) (Figure 3D).

**Figure 3.**
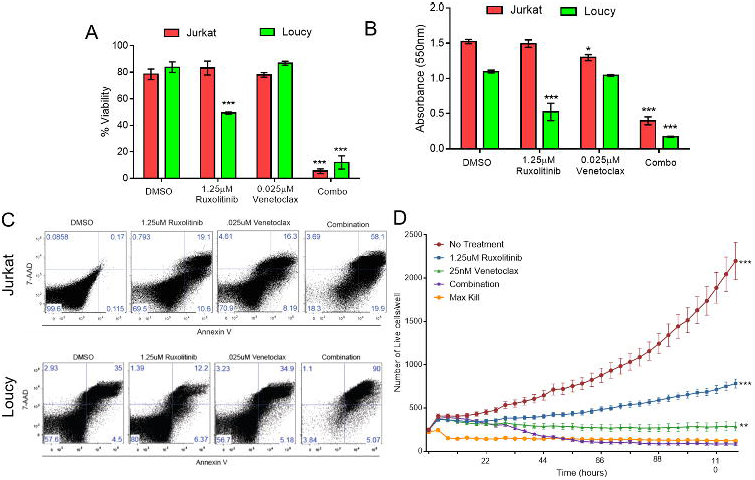
Ruxolitinib and venetoclax combination treatment is more potent than single-agent treatment. Trypan blue (A), MTT proliferation (B), Annexin V/7-AAD flow cytometric (C) and IncuCyte live cell imaging (D) data of single agent and combination ruxolitinib and venetoclax treatment of the indicated doses at 48 hours compared to a DMSO control. Two independent experiments were performed with triplicate measurements. * p < 0.05; ** p < 0.01; *** p < 0.001.

### Ruxolitinib and venetoclax fail to treat T-ALL *in vivo* due to leukemia CNS infiltration

Based on the efficacy of ruxolitinib and venetoclax observed *in vitro*, a xenograft model of T-ALL was established utilizing Jurkat cells and NSG mice[28,29] to test these drugs as a combination therapy *in vivo*. While single agent studies have used venetoclax doses up to 100mg/kg/day[12], we observed that these doses were not well tolerated when given in combination with 30 mg/kg/day ruxolitinib causing lethal gastrointestinal toxicity in NSG mice minutes to hours after administration. However, we did find that 35 mg/kg/day venetoclax was able to be tolerated, and so was used in our combination treatment groups with 30mg/kg/day ruxolitinib (and as a single agent dose as a control) for 14 days after cancer burden establishment. No diarrhea or other gastrointestinal toxicity was observed. *In vivo* dosing revealed optimal doses of 30 mg/kg/day of ruxolitinib and 35 mg/kg/day of venetoclax delivered via oral gavage once daily[15,17,30,31] for 14 days after cancer burden establishment (Figure 4A). Surprisingly, ruxolitinib and venetoclax failed to improve clinical scores (Figure 4B) or survival (Figure 4C) after T-ALL challenge when given as single agents or in combination with the cause of death ultimately due to hind limb paralysis (Supplemental Figure 4), suggesting using reduced doses to limit toxicity may have reduced efficacy.

**Figure 4.**
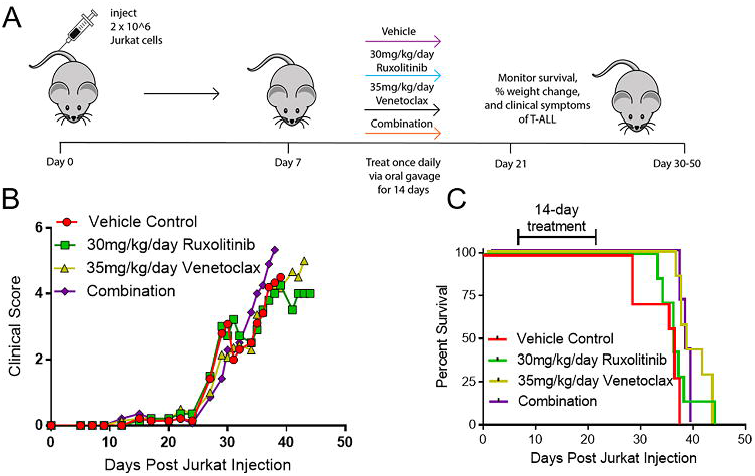
Ruxolitinib and venetoclax single agent and combination treatment is not efficacious against T-ALL in vivo. (A) Schematic of experimental design. (B-C) Clinical score (B) and percent survival (C) of single agent ruxolitinib, venetoclax or combination treatment given by oral gavage once daily for 14 days starting at seven days post Jurkat transplant. N=5 mice/group.

We next injected NSG mice with GFP expressing Jurkat cells and analyzed the cancer burden (without ruxolitinib or venetoclax treatment) by flow cytometry for an anti-human CD45^+^GFP^+^ double population in the bone marrow, spleen, brain, and spinal cord over the course of the disease (approximately 50-60 days). Interestingly, the T-ALL burden was primarily in the spinal cord and eventually the brain, but not the bone marrow and spleen (Figure 5A-C). While both inhibitors are FDA approved and currently used in the clinic, there is no published data that shows if either inhibitor has the ability to effectively penetrate the CNS and treat T-ALL. Naïve mice were treated by oral gavage with the highest published *in vivo* single dose of either ruxolitinib (50 mg/kg)[30] or venetoclax (100 mg/kg),[15] and were then sacrificed at 3 hours post-ruxolitinib or 16 hours post-venetoclax[32,33]. The peripheral blood, spinal cord, and brain were then analyzed for the presence of either inhibitor by LC-MS-MS. We observed that there was a 5X and 100X reduction in the amount of ruxolitinib and venetoclax respectively in the spinal cord/brain compared to the serum (Figure 5D). These data show that when administered *in vivo* at doses of 30 mg/kg/day ruxolitinib and 35 mg/kg/day venetoclax, these drugs insufficiently penetrate the BBB, potentially explaining why they fail to treat CNS T-ALL disease burden in the Jurkat *in vivo* model.

**Figure 5.**
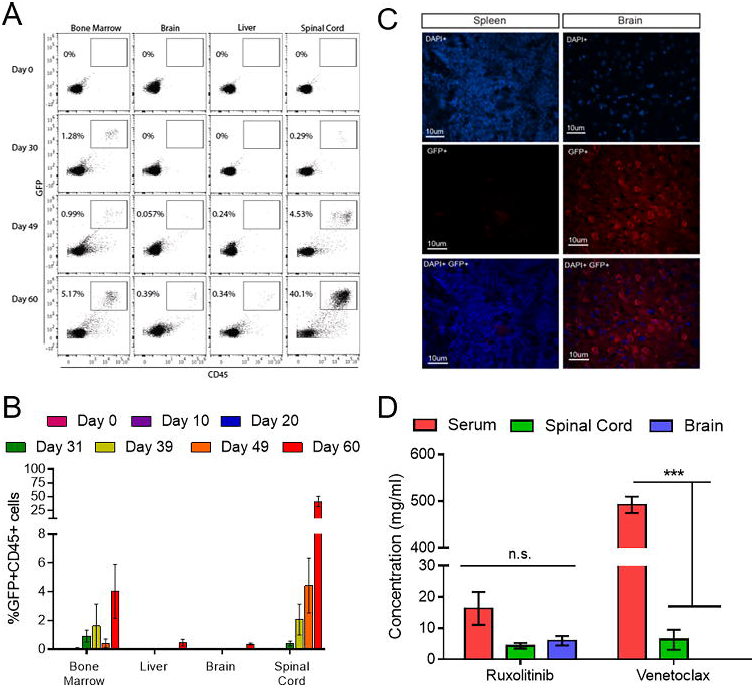
T-ALL evades ruxolitinib and venetoclax treatment in the CNS. (A-B) Flow cytometric analysis of human CD45^+^/GFP^+^ Jurkat cells in the bone marrow, brain, liver and spinal cord of NSG mice between 10 and 60 days post-transplant. (C) Immunohistochemistry of spleen and brain sections stained with DAPI (blue) and an anti-GFP/AF-555 secondary antibody (red). (D) LC-MS-MS analysis of ruxolitinib and venetoclax concentration in the serum, spinal cord and brain of NSG mice after an oral gavage of 50mg/kg of ruxolitinib or 100mg/kg of venetoclax. N= 3 mice at each time point. * p < 0.05; ** p < 0.01; *** p < 0.001.

### The CXCR4/CXCL12 pathway is upregulated in Jurkat cells and mediates CNS penetration

To determine how Jurkat cells were infiltrating the CNS, we first examined the CXCR4 expression on Jurkat cells compared to healthy human controls. The chemokine CXCR4 recognizes CXCL12 or stromal-derived factor 1 (SDF-1) that directs cells to the CNS. We found that Jurkat cells has significantly elevated levels of CXCR4 compared to healthy human CD3^+^ cells (Figure 6A).

**Figure 6.**
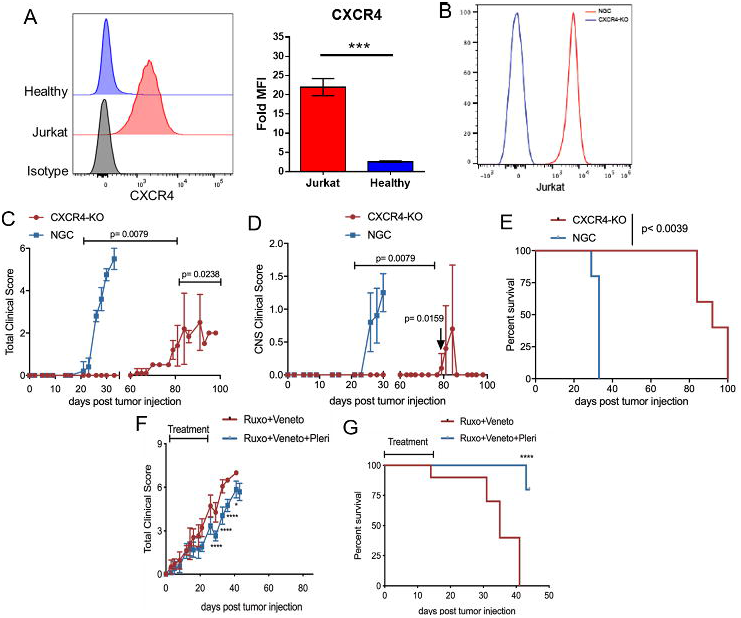
Blocking CXCR4 prevents CNS infiltration and facilitates combination therapy efficacy. (A) Histogram of CXCR4 expression on Jurkat cells and CD3^+^ healthy adult T cells compared to an isotype control. Bar graph represents two independent experiments. (B) Histogram of successful CXCR4 expression reduction in Jurkat cells (blue line) compared to a no guide control (red line). (C-E) Clinical score (C), CNS clinical score according to Supplemental Table 1 (D) and overall survival (E) for control Jurkat cells (blue) and CXCR4-KO Jurkat cells (red). (F-G) Clinical score (F) and overall survival (G) are shown for mice injected with wild-type Jurkat cells and treated with either ruxolitinib/venetoclax (red line) or ruxolitinib/venetoclax and the CXCR4 inhibitor plerixafor (blue line). N=3-10 mice/group * p < 0.05; ** p < 0.01; *** p < 0.001.

Using CRISPR-Cas9 technology we deleted the CXCR4 gene in Jurkat cells (Figure 6B). We then injected the CXCR4-KO, or no-guide control (NGC) Jurkat cells, into mice and monitored them for clinical symptoms of T-ALL. The NGC group developed rapid symptoms of CNS disease and had to be euthanized by day 35 while the mice transplanted with CXCR4-KO cells survived until day 80-100 and only had a few mice develop signs of CNS pathology (Figure 6C-E). Lastly, in combination with ruxolitinib and venetoclax therapy, we used two different approaches to block CNS infiltration in vivo that are relevant to the clinic. The first approach attempted to block CXCR4 with an anti-CXCR4 antibody but no efficacy was detected (Supplemental Figure 5). The second approach used plerixafor, a small molecule inhibitor of the CXCR4/CXCL12 axis. The use of plerixafor in combination with ruxolitinib and venetoclax significantly reduced clinical scores (Figure 6F) and extended survival (Figure 6G) of mice compared to the condition without plerixafor. Overall, these data show that blocking T-ALL infiltration and persistence in the CNS with a CXCR4 small molecule inhibitor is a feasible strategy to prevent T-ALL escape from treatment with ruxolitinib and venetoclax.

## Discussion

Relapsed or refractory T-ALL remains a significant challenge to manage, especially since most patients are heavily pretreated with chemotherapy that causes significant end organ toxicity. Development of targeted therapies that inhibit pathways exploited by T-ALL to augment proliferation and survival may improve outcomes while also minimizing systemic toxicity. Prior studies involving RNA or whole genome sequencing of T-ALL samples identified the JAK/STAT pathway[26,34,35] and the BCL2 pathway as being dysregulated[15]. In this study ruxolitinib, a JAK1/2 inhibitor, showed efficacy against T-ALL *in vitro* as measured by viability and proliferation assays. We also found that venetoclax, a potent BCL2 inhibitor demonstrated the same effects *in vitro*. Furthermore, it was demonstrated that when employing these two inhibitors together *in vitro*, the combination therapy demonstrated an increased effect as measured by increased apoptosis and a decrease in proliferation compared to either inhibitor alone. This data confirmed findings previously demonstrated in T-ALL[20,36,37] and other cancers.[38-40]

Ruxolitinib treatment alone has been considered a breakthrough therapy for myeloproliferative neoplasms but long-term studies have shown that this treatment is not curative and does not lead to molecular or pathological remission. This may be caused by the disease evading treatment and utilizing an alternative survival pathway, such as the BCL2 pathway. While Senkevitch et al 2018 demonstrated a robust synergistic effect of combined JAK/STAT and BCL2 inhibition *in vivo* in IL7R*α*+ T-ALL, all mice succumbed to T-ALL[20]. Our study demonstrates for the first time that T-ALL progression after combined ruxolitinib and venetoclax treatment arises from CNS invasion mediated by CXCR4, a homing pathway previously identified in untreated T-ALL cells.[22] The xenograft model of T-ALL with Jurkat cells presents as CNS disease with the cancer burden homing primarily into the spinal cord, rather than a primary systemic form where the burden is primarily seen in the bone marrow, spleen, and liver.[41] This xenograft model has clinical significance given that CNS relapse is common for T-ALL, and is a strong driver for a poor prognosis.[42]

The current standard of treating CNS leukemia is either by administering repeated doses of intrathecal chemotherapy or giving irradiation to the neuro-axis. Identification of agents that are effective in the CNS when administered systemically is an urgent need. This study demonstrated for the first time that while ruxolitinib and venetoclax have activity *in vitro*, they may be ineffective *in vivo* since they are unable to effectively cross the BBB and target CNS leukemia. LC-MS-MS demonstrated that low amounts of ruxolitinib (administered at the highest published dose 50mg/kg)[30] can be detected in the spinal cord and brain of NSG mice and low amounts of venetoclax (administered at the highest published dose 100mg/kg[1,15]) can be detected in the spinal cord, though they are not high enough concentrations to be expected to have an anti-tumor effect at a physiological dose (ruxolitinib 30mg/kg/day; venetoclax 35mg/kg/day). These data could be informative as future generations of these small molecule inhibitors may need to be engineered to allow for better CNS penetration.

Further we dissected the homing mechanism demonstrated with Jurkat cells in NSG mice. Several studies of human T-ALL have demonstrated that CNS relapsed T-ALL occurs in part through the CXCR4-CXCL12 chemokine pathway[22,24,25,43]. We confirmed that Jurkat cells utilize this pathway to escape treatment by ruxolitinib and venetoclax and infiltrate the CNS. CXCR4 is abundantly present on the surface of Jurkat cells and CXCL12 is highly abundant in the CNS. Since murine CXCL12 can bind the human CXCR4 receptor, human T-ALL can enter the CNS of NSG mice[44,45]. Utilizing CRISPR-Cas9 to delete CXCR4 from the surface of Jurkat cells, we determined that CXCR4 plays a critical role in Jurkat cell homing to not only the CNS but also non-CNS sites. CXCR4 deletion or inhibition in T-ALL cells significantly delays lethality by 60-70 days while reducing overall and neurological clinical scores, showing that CXCR4 plays a critical role in T-ALL progression. Interestingly though, the use of an anti-CXCR4 antibody failed to prevent CNS penetration. This is most likely due to the poor ability of antibodies to cross the BBB with one report suggesting antibodies exist at 1000X lower concentration in the CNS than in the serum[46]. Additionally, we treated our mice with anti-CXCR4 after Jurkat injection suggesting that Jurkat cells had already infiltrated the CNS and evaded anti-CXCR4 treatment. Since plerixafor was efficacious in our model, we hypothesize that plerixafor is capable of crossing the BBB, blocking CXCR4 signaling and mediating the egress of Jurkat cells out of the CNS.

Development of novel therapies for relapsed/refractory T-ALL is a high unmet need given the poor prognosis of this cancer in children and adults. Inhibiting the JAK/STAT pathway along with BCL2, using ruxolitinib and venetoclax, is an effective treatment for relapsed T-ALL, but requires additional inhibition of CXCR4 to prevent CNS invasion and relapse after treatment. Future studies could consider targeting all three pathways as a means of inducing a sustained complete remission. Combination of ruxolitinib and venetoclax with cell-based therapies, like T or NK cells, could also be considered since immune effector cells can easily cross the BBB, but the impact of these drugs on these cell subsets will need to be explored carefully.

## Acknowledgements

This work was supported by grants from the NHLBI/NIH T32 HL07899 (K.L.W.), NIH TL1 TR002375 (S.P.R), Cormac Pediatric Leukemia Postdoctoral Fellowship (N.J.H.), American Association for Immunologists Careers in Immunology Fellowship (M.N.B.), NCI/NIH T32 CA009135 (M.M.C.), NSF 1810916 WiscAMP Bridge to the Doctorate and NSF Graduate Research Fellowship Program DGE-1747503 (A.E.Q.), American Society of Hematology HONORS award (T.B.G.), NCI/NIH R01 CA187299 (L.R.), St. Baldrick’s-Stand Up To Cancer Pediatric Dream Team Translational Research Grant SU2C-AACR-DT-27-17, Vince Lombardi Cancer Foundation, NCI/NIH K08 CA174750, NCI/NIH R01 CA215461 and the MACC Fund (C.M.C). We would like to thank Nicole Piscopo and Krishanu Saha for providing the Jurkat-GFP cell line. We would like to thank the UWCCC Flow Cytometry core facility and UWCCC Experimental Pathology core facility, who are supported in part through NCI/NIH P30 CA014520 as well as the UW Biotechnology Center Genome Editing and Animal Model core facility as well as the Mass Spectrometry core facility, who is supported in part through NIGMS/NIH P50 GM64598 and NIDDK/NIH R33 DK070297 and the National Science Foundation (DBI-0520825, DBI-9977525). Stand Up To Cancer is a division of the Entertainment Industry Foundation.

Research grants are administered by the American Association for Cancer Research, the scientific partner of SU2C. Any opinions, findings, and conclusions or recommendations expressed in this material are those of the author(s) and do not necessarily reflect the views of the National Science Foundation or the Department of Health and Human Services, nor does mention of trade names, commercial products, or organizations imply endorsement by the US Government. None of these funding sources had any input in the study design, analysis, manuscript preparation or decision to submit for publication.

## Authorship Contributions

K.L.W. and C.M.C designed the experiments, analyzed and interpreted results and wrote the manuscript; N.J.H, S.P.R., S.A.K, F.Z., S.L.O, M.M.C., A.E.Q., A.S.F., and T.B.G. conducted experiments and analyzed data; N.J.H, S.A.K. and S.L.O analyzed data and generated figures; and S.P.R., D.P.T., M.N.B. and L.R. discussed and interpreted results. All authors read and approved the manuscript.

## References

1. Sanda T, Tyner JW, Gutierrez A, et al. TYK2-STAT1-BCL2 pathway dependence in T-cell acute lymphoblastic leukemia. Cancer Discov. 2013 May;3(5):564–77.

2. Follini E, Marchesini M, Roti G. Strategies to Overcome Resistance Mechanisms in T-Cell Acute Lymphoblastic Leukemia. Int J Mol Sci. 2019 Jun 20;20(12).

3. Marks DI, Rowntree C. Management of adults with T-cell lymphoblastic leukemia. Blood. 2017 Mar 2;129(9):1134–1142.

4. Asselin BL, Devidas M, Wang C, et al. Effectiveness of high-dose methotrexate in T-cell lymphoblastic leukemia and advanced-stage lymphoblastic lymphoma: a randomized study by the Children’s Oncology Group (POG 9404). Blood. 2011 Jul 28;118(4):874–83.

5. Van Vlierberghe P, Ferrando A. The molecular basis of T cell acute lymphoblastic leukemia. J Clin Invest. 2012 Oct;122(10):3398–406.

6. Goldberg JM, Silverman LB, Levy DE, et al. Childhood T-cell acute lymphoblastic leukemia: the Dana-Farber Cancer Institute acute lymphoblastic leukemia consortium experience. J Clin Oncol. 2003 Oct 1;21(19):3616–22.

7. Marks DI, Paietta EM, Moorman AV, et al. T-cell acute lymphoblastic leukemia in adults: clinical features, immunophenotype, cytogenetics, and outcome from the large randomized prospective trial (UKALL XII/ECOG 2993). Blood. 2009 Dec 10;114(25):5136–45.

8. Kucine N, Marubayashi S, Bhagwat N, et al. Tumor-specific HSP90 inhibition as a therapeutic approach in JAK-mutant acute lymphoblastic leukemias. Blood. 2015 Nov 26;126(22):2479–83.

9. Delgado-Martin C, Meyer LK, Huang BJ, et al. JAK/STAT pathway inhibition overcomes IL7-induced glucocorticoid resistance in a subset of human T-cell acute lymphoblastic leukemias. Leukemia. 2017 Dec;31(12):2568–2576.

10. Liszewski W, Naym DG, Biskup E, et al. Psoralen with ultraviolet A-induced apoptosis of cutaneous lymphoma cell lines is augmented by type I interferons via the JAK1-STAT1 pathway. Photodermatol Photoimmunol Photomed. 2017 May;33(3):164–171.

11. Senkevitch E, Durum S. The promise of Janus kinase inhibitors in the treatment of hematological malignancies. Cytokine. 2017 Oct;98:33–41.

12. Messina NL, Banks KM, Vidacs E, et al. Modulation of antitumour immune responses by intratumoural Stat1 expression. Immunol Cell Biol. 2013 Oct;91(9):556–67.

13. Seymour JF, Kipps TJ, Eichhorst B, et al. Venetoclax-Rituximab in Relapsed or Refractory Chronic Lymphocytic Leukemia. N Engl J Med. 2018 Mar 22;378(12):1107–1120.

14. Seymour JF, Mobasher M, Kater AP. Venetoclax-Rituximab in Chronic Lymphocytic Leukemia. N Engl J Med. 2018 May 31;378(22):2143–2144.

15. Chonghaile TN, Roderick JE, Glenfield C, et al. Maturation stage of T-cell acute lymphoblastic leukemia determines BCL-2 versus BCL-XL dependence and sensitivity to ABT-199. Cancer Discov. 2014 Sep;4(9):1074–87.

16. Koppikar P, Bhagwat N, Kilpivaara O, et al. Heterodimeric JAK-STAT activation as a mechanism of persistence to JAK2 inhibitor therapy. Nature. 2012 Sep 6;489(7414):155–9.

17. Peirs S, Matthijssens F, Goossens S, et al. ABT-199 mediated inhibition of BCL-2 as a novel therapeutic strategy in T-cell acute lymphoblastic leukemia. Blood. 2014 Dec 11;124(25):3738–47.

18. Matulis SM, Gupta VA, Nooka AK, et al. Dexamethasone treatment promotes Bcl-2 dependence in multiple myeloma resulting in sensitivity to venetoclax. Leukemia. 2016 May;30(5):1086–93.

19. Zhang M, Mathews Griner LA, Ju W, et al. Selective targeting of JAK/STAT signaling is potentiated by Bcl-xL blockade in IL-2-dependent adult T-cell leukemia. Proc Natl Acad Sci U S A. 2015 Oct 6;112(40):12480–5.

20. Senkevitch E, Li W, Hixon JA, et al. Inhibiting Janus Kinase 1 and BCL-2 to treat T cell acute lymphoblastic leukemia with IL7-Ralpha mutations. Oncotarget. 2018 Apr 27;9(32):22605–22617.

21. Pui CH, Robison LL, Look AT. Acute lymphoblastic leukaemia. Lancet. 2008 Mar 22;371(9617):1030–43.

22. T RJ. Role of CXCR4-mediated bone marrow colonization in CNS infiltration by T cell acute lymphoblastic leukemia [Article]. Journal of Leukocyte Biology. 2016;99:1077–1087.

23. Pitt LA, Tikhonova AN, Hu H, et al. CXCL12-Producing Vascular Endothelial Niches Control Acute T Cell Leukemia Maintenance. Cancer Cell. 2015 Jun 8;27(6):755–68.

24. Passaro D, Irigoyen M, Catherinet C, et al. CXCR4 Is Required for Leukemia-Initiating Cell Activity in T Cell Acute Lymphoblastic Leukemia. Cancer Cell. 2015 Jun 8;27(6):769–79.

25. Yao H, Price TT, Cantelli G, et al. Leukaemia hijacks a neural mechanism to invade the central nervous system. Nature. 2018 Aug;560(7716):55–60.

26. Liu Y, Easton J, Shao Y, et al. The genomic landscape of pediatric and young adult T-lineage acute lymphoblastic leukemia. Nat Genet. 2017 Aug;49(8):1211–1218.

27. Anderson NM, Harrold I, Mansour MR, et al. BCL2-specific inhibitor ABT-199 synergizes strongly with cytarabine against the early immature LOUCY cell line but not more-differentiated T-ALL cell lines. Leukemia. 2014 May;28(5):1145–8.

28. Maude SL, Tasian SK, Vincent T, et al. Targeting JAK1/2 and mTOR in murine xenograft models of Ph-like acute lymphoblastic leukemia. Blood. 2012 Oct 25;120(17):3510–8.

29. Wu X, Zhang LS, Toombs J, et al. Extra-mitochondrial prosurvival BCL-2 proteins regulate gene transcription by inhibiting the SUFU tumour suppressor. Nat Cell Biol. 2017 Oct;19(10):1226–1236.

30. Carniti C, Gimondi S, Vendramin A, et al. Pharmacologic Inhibition of JAK1/JAK2 Signaling Reduces Experimental Murine Acute GVHD While Preserving GVT Effects. Clin Cancer Res. 2015 Aug 15;21(16):3740–9.

31. Khaw SL, Suryani S, Evans K, et al. Venetoclax responses of pediatric ALL xenografts reveal sensitivity of MLL-rearranged leukemia. Blood. 2016 Sep 8;128(10):1382–95.

32. Haile WB, Gavegnano C, Tao S, et al. The Janus kinase inhibitor ruxolitinib reduces HIV replication in human macrophages and ameliorates HIV encephalitis in a murine model. Neurobiol Dis. 2016 Aug;92(Pt B):137–43.

33. Eisenmann ED, Jin Y, Weber RH, et al. Development and validation of a sensitive UHPLC-MS/MS analytical method for venetoclax in mouse plasma, and its application to pharmacokinetic studies. J Chromatogr B Analyt Technol Biomed Life Sci. 2020 May 20;1152:122176.

34. Zhang J, Ding L, Holmfeldt L, et al. The genetic basis of early T-cell precursor acute lymphoblastic leukaemia. Nature. 2012 Jan 11;481(7380):157–63.

35. Gianfelici V, Chiaretti S, Demeyer S, et al. RNA sequencing unravels the genetics of refractory/relapsed T-cell acute lymphoblastic leukemia. Prognostic and therapeutic implications. Haematologica. 2016 Aug;101(8):941–50.

36. Degryse S, de Bock CE, Demeyer S, et al. Mutant JAK3 phosphoproteomic profiling predicts synergism between JAK3 inhibitors and MEK/BCL2 inhibitors for the treatment of T-cell acute lymphoblastic leukemia. Leukemia. 2018 Mar;32(3):788–800.

37. Kontro M, Kuusanmaki H, Eldfors S, et al. Novel activating STAT5B mutations as putative drivers of T-cell acute lymphoblastic leukemia. Leukemia. 2014 Aug;28(8):1738–42.

38. Karjalainen R, Pemovska T, Popa M, et al. JAK1/2 and BCL2 inhibitors synergize to counteract bone marrow stromal cell-induced protection of AML. Blood. 2017 Aug 10;130(6):789–802.

39. Kuusanmaki H, Leppa AM, Polonen P, et al. Phenotype-based drug screening reveals association between venetoclax response and differentiation stage in acute myeloid leukemia. Haematologica. 2019 Jul 11.

40. Waibel M, Solomon VS, Knight DA, et al. Combined targeting of JAK2 and Bcl-2/Bcl-xL to cure mutant JAK2-driven malignancies and overcome acquired resistance to JAK2 inhibitors. Cell Rep. 2013 Nov 27;5(4):1047–59.

41. Vadillo E, Dorantes-Acosta E, Pelayo R, et al. T cell acute lymphoblastic leukemia (T-ALL): New insights into the cellular origins and infiltration mechanisms common and unique among hematologic malignancies. Blood Rev. 2018 Jan;32(1):36–51.

42. Nguyen K, Devidas M, Cheng SC, et al. Factors influencing survival after relapse from acute lymphoblastic leukemia: a Children’s Oncology Group study. Leukemia. 2008 Dec;22(12):2142–50.

43. Burger JA, Peled A. CXCR4 antagonists: targeting the microenvironment in leukemia and other cancers. Leukemia. 2009 Jan;23(1):43–52.

44. Beider K, Nagler A, Wald O, et al. Involvement of CXCR4 and IL-2 in the homing and retention of human NK and NK T cells to the bone marrow and spleen of NOD/SCID mice. Blood. 2003 Sep 15;102(6):1951–8.

45. Hess NJ, Lindner PN, Vazquez J, et al. Different Human Immune Lineage Compositions Are Generated in Non-Conditioned NBSGW Mice Depending on HSPC Source. Front Immunol. 2020;11:573406.

46. Neves V, Aires-da-Silva F, S C-R, et al. Antibody Approaches to Treat Brain Diseases. Trends in Biotechnology. 2016;34(34):36–48.

