## Supplemental Figure Legends and Methods for "CXCR4 allows T cell acute lymphoblastic leukemia to escape from JAK1/2 and BCL2 inhibition through CNS infiltration"

Supplementary Figure 1. Loucy early precursor-ALL can be targeted by ruxolitinib and venetoclax (A) Trypan blue exclusion data of cells treated with serial diluted ruxolitinib at 24, 48 and 72 h (B) Trypan blue exclusion data of cells treated with serial diluted venetoclax at 24, 48, 72 h (C) MTT proliferation data of cells treated with serial diluted ruxolitinib at 24, 48, and 72 h (D) MTT proliferation data of cells treated with serial diluted venetoclax at 24, 48, 72 h. Experiments were duplicated with (n =3) each experiment.

Supplementary Figure 2. Ruxolitinib and venetoclax decrease survival and proliferation of Loucy cells (A) Trypan blue exclusion data of combination treatment and single dose controls (B) MTT proliferation data of combination treatment and single dose controls. (C) Annexin V and & 7-AAD data of combination treatment and single dose controls after 48 hour treatment. Experiments were duplicated with (n =3) each experiment.

Supplementary Figure 3. Target analysis of ruxolitinib and venetoclax (A) Western blot analysis of JAK/STAT and BCL2 proteins in Jurkat cells treated with vehicle control, ruxolitinib, venetoclax or combination for 24h. Human NK cell lysate was used a positive (+) control. (B) Statistical representation of the fold change of protein density normalized to positive control compared to DMSO control (C) Flow analysis of pSTAT1 and pSTAT3.

Supplementary Figure 4. Single agent therapy with ruxolitinib or venetoclax does not improve survival after Jurkat leukemia challenge. NSG mice were challenged with Jurkat cells on Day +0, and once established leukemia was measured in peripheral blood (>1%) mice were treated with (A) increasing doses of ruxolitinib (Ruxo) vs. vehicle control (VC), or (B) increasing doses of venetoclax (Veneto) vs. vehicle control (VC) for a 21 day course. Overall survival was measured. N = 3 mice/group

Supplementary Figure 5. Addition of anti-CXCR4 monoclonal antibody to ruxolitinib and venetoclax combination therapy does not improve survival. NSG mice were challenged with Jurkat cells on Day +0, and then mice were treated with 30mg/kg/day ruxolitinib and 35mg/kg/day venetoclax for 21 days. Mice were randomized to receive 10ug anti-CXCR4 or isotype control for 2 days starting on Day +7 and then again on Day +21. All mice were followed for overall survival, total clinical scores and CNS clinical scores. N = 5 mice/group. * = p < 0.05

**Supplementary Materials and Methods**

*Organ harvesting and processing:*

Bone marrow, spleen, liver, spinal cord, and brain were harvested from mice and processed into single cell suspensions. Brain was digested with 2 ml of 2 mg/ml collagenase D (Cat # LS004188, Worthington Biochemical, Lakewood, NJ) + 0.2% BSA + 28 U/ml DNAse I (Cat # 10 104 159 001, Roche, Basel, Switzerland) in 1x HBSS for 30 min at 37 °C. In order to measure levels of ruxolitinib in brain and spinal cord, mice underwent cardiac perfusion with 1x PBS prior to organ extraction. Brain and spinal cord were snap frozen and stored at -80 °C until homogenized into a powder where it was then further processed for LC-MS-MS. Homogenate of each organ and serum was extracted with MeOH + 10% H_2_O.

*Viability and proliferation assays:*

Trypan blue exclusion assays used a 1:1 ratio of cells to trypan blue. MTT Proliferation Assays (Cat # 11465007001, Roche, Basel, Switzerland) were done following the manufacturer’s instructions. Serial dilution of DMSO, ruxolitinib (5 μM- 0.157μM), and venetoclax (0.05 μM - 0.0016 μM) were added to wells of Jurkat-GFP cells in triplicate. The absorption of the MTT proliferation assay were read on a SpectraMax Plus (Molecular Devices, San Jose, CA) at 550nm. Live cell imaging analysis was conducted via the Incucyte (Sartorius, Göttingen, Germany) for 5 days. Viability was confirmed using the GFP expression endogenous to the Jurkat-GFP expressing cell line. Max kill group was plated with 1% Triton.

*Flow cytometry analysis:*

All flow cytometry antibodies were purchased from Biolegend, San Diego, CA unless otherwise stated. Jurkat cells were stained with Annexin V - APC (Cat # 640941) and 7-AAD (Cat # 420404) after incubation with inhibitors. Jurkat cells were also stained with CD184/CXCR4 - PE/Cy5.5 (Cat # 306508) and Ghost Dye Red 780 (Cat # 13-0865-T500, Tonbo Biosciences, San Diego, CA). Single cell suspensions of organs were stained with anti-human CD45 - BV510 (Cat # 304036) or PE (Cat # 368510) and DAPI (Cat # 422801).

*Immunofluorescence analysis:*

Immunofluorescence was performed on the spleen and brain of mice throughout the experiment. Organs were cryo-embedded in OCT solution (Sakura Finetek, Torrance, CA) and sectioned. Frozen sections were cut at 10 μ, mounted on slides, air dried, fixed in 4 °C acetone, then air dried. Slides were rinsed in dH_2_O for 10 min, blocked for 1 h in 10% goat serum in 1x PBS at 20 °C, and labeled with 1:200 anti-GFP (clone D5.1, Cat # 2956, Cell Signaling Technology, Danvers, MA) In 1x PBS with 1% goat serum overnight at 4 °C. Slides were then stained with 1:1000 goat anti-rabbit IgG Alexafluor 555 in 1x PBS (Cat # A21428, Invitrogen, Carlsbad, CA) for 30 min at 20 °C in the dark. Slides were then stained with Prolong Gold antifade reagent (ThermoFisher, Waltham, MA) with DAPI.

*Western blot analysis*

Cells were collected and lysed using MAPK lysis buffer (4 mM sodium pyrophosphate, 50 mM HEPES, 100 mM NaCl, 1 mM EDTA, 10 mM NaF, 2 mM orthovanadate, pH 7.5) with protease inhibitor cocktail (Cat # P8340, Sigma Aldrich, St. Louis, MO; ThermoFisher, Waltham, MA) Protein concentrations were determined by BCA Assay (Cat # 23250, Thermo Fisher, Waltham, MA). Proteins were separated by SDS-PAGE and transferred to a nitrocellulose membrane. Membrane was blocked for 30 min with 5% non-fat milk in TBST at 20 °C, and incubated overnight at 4 °C with the following antibodies from Cell Signaling Technology, Danvers, MA unless otherwise stated: STAT1 (Cat #9172L), pSTAT1 (Tyr701), (Cat #7649S), STAT3 (Cat # 124H6), pSTAT3 (Tyr705), (Cat #9145), STAT5 (Cat #25656S), pSTAT5 (Tyr695) (Cat #9314S), BCL2 (Santa Cruz Biotechnology, sc7382), BIM (Cat #2933S), BAX (Cat #2774S), BAK (Cat #121058), Cytochrome C (Cat # 11940S), H3 (abcam, ab1791), β-Actin (Cat #4967), and CXCL12 antibody (Cat #3740). Membranes were then incubated for 1 h with 1:5000 anti-rabbit (Cat# #7074) or anti-mouse (Cat #7076) HRP- conjugated secondary and developed using the chemiluminescence HRP substrate kit (Cat # 34080, 34095, Thermo Fisher, Waltham, MA).

CRISPR-Cas9 deletion of CXCR4 in T-ALL

CXCR4 deletion was performed at the Genome Editing and Animal Models Core at University of Wisconsin-Madison. Briefly, Jurkat cells were cultured in RPMI-1640 (Gibco, Waltham. MA) containing 10% FBS (ATCC), 10 U/mL Penicillin, 10 μg/ml Streptomycin, and 25 ng/mL Amphotericin B (ATCC). RNP delivery occurred via electroporation. All CRISPR reagents were purchased from IDT, Coralville, IA. RNP complex was prepared immediately before electroporation. 2x10^5^ cells were used for each electroporation. CXCR4 exon 2 CRISPR RNA (crRNA) (TACACCGAGGAAATGGGCTCAGG) and Atto-trans-activating crRNA (Atto-tracrRNA) were mixed in equimolar concentrations (200 μM) to form a ctRNA::tracrRNA complex (ctRNA). The molar ratio ctRNA to Cas9 was 1:1 with the working concentration of the electroporation being 2 μM Cas9:ctRNA and 2 μM Alt-R Cas9 electroporation enhancer. RNP was delivered into cells using a Lonza (Basil, Switzerland) 4D-X Nucleofector according to manufacturer’s specifications. Single cell sorting occurred 40 h after electroporation. The cells were harvested, washed, and treated with APC anti-human CXCR4 antibody (Cat #306509, Biolegend, San Diego, CA) and DAPI in accordance with the manufacturer’s specifications. Cells were then put on a BD FACSAria (BD Biosciences, Franklin Lakes, NJ) and sorted for APC^-^Atto^+^ population into 1:1 conditioned growth media (fresh media: filtered media from fully confluent flask) for clonal outgrowth. Surviving clones were genotyped via amplicon sequencing on the MiSeq. The targeted region from each was then amplified via PCR, barcoded with a second PCR reaction, pooled, and run on a MiSeq Nano (University of Wisconsin DNA Sequencing Facility, Madison, WI). Data analysis was performed with CRISPResso (PMID: 27404874). Selected clones carried compound heterozygous nonsense mutations in the targeted region.

*Liquid chromatography and tandem mass spectrometry:*

Analyses of ruxolitinib and venetoclax in organs were carried out on the LC/MS/MS Linear Ion Trap Quadrupole system consisting of an Agilent 1100 High Pressure Liquid Chromatograph (Santa Clara, CA) placed on the front end of an Applied Biosystems 3200 Q-Trap Mass Spectrometer equipped with a Turbo V™ (Beverly Hills, CA) spray source running in the Multiple Reaction Monitoring mode. Chromatography separations were accomplished using Agilent ZORBAX SB-C18 Rapid Resolution HT reversed-phase column 2.1x 50 mm (1.8 µm particle size) onto which 1 µl of methanol extracted sample was automatically loaded. HPLC delivered solvents A: 0.1% (v/v) formic acid in water, and B: 0.1% formic acid in acetonitrile at 0.2 ml/min. After 2 min loading and equilibration at 95% buffer A, the analytes were eluted from the column directly into the electrospray orifice over 23 min 5% (v/v) B to 100% (v/v) B fast gradient, followed by a 5 min hold at 100% (v/v) B and 9min post gradient equilibration at 95% buffer A. As analytes eluted from HPLC-column into the electrospray source MRM mode selectively measured six transitions based on precursor mass and specific fragment mass for drugs of interest. The transitions measured were: m/z= 306.4, fragment m/z= 186.1 for most abundant MS/MS fragment of ruxolitinib; m/z= 306.4, fragment m/z= 159.0 for second most abundant MS/MS fragment of ruxolitinib; m/z= 435.0, fragment m/z= 321.1 for most abundant MS/MS fragment of doubly charged [M+2H]^2+^ venetoclax; m/z= 435.0, fragment m/z= 163.4 for second most abundant MS/MS fragment of doubly charged [M+2H]^2+^ venetoclax; m/z= 868.5, fragment m/z= 321.2 for most abundant MS/MS fragment of singly charged [M+H]^+^ venetoclax and m/z= 868.5, fragment m/z= 233.1 for second most abundant MS/MS fragment of singly charged [M+H]^+^ venetoclax. The following instrumental parameters were used to generate the most optimum protonated ions and selectively diagnostic fragment ions: Ion spray voltage (IS), 4500 V; Curtain gas (CUR), 25 psi; Nebulizer gas (GS1), 50 psi; Turbo gas (GS2), 30 psi; Turbo gas temperature (TEM), 650 °C; Interface heather (ihe), ON; Collision Gas (CAD), Medium; Declustering potential (DP), 50 eV; Entrance potential (EP), 10 eV; Collision energy (CE), 40 eV; Collision cell exit potential (CXP), 5 eV; Detector (CEM), 2500; Dwell time, 200 msec with pause time, 5 msec.
