## Supplementary Figures 1-5 and Table 1 for "CXCR4 allows T cell acute lymphoblastic leukemia to escape from JAK1/2 and BCL2 inhibition through CNS infiltration"

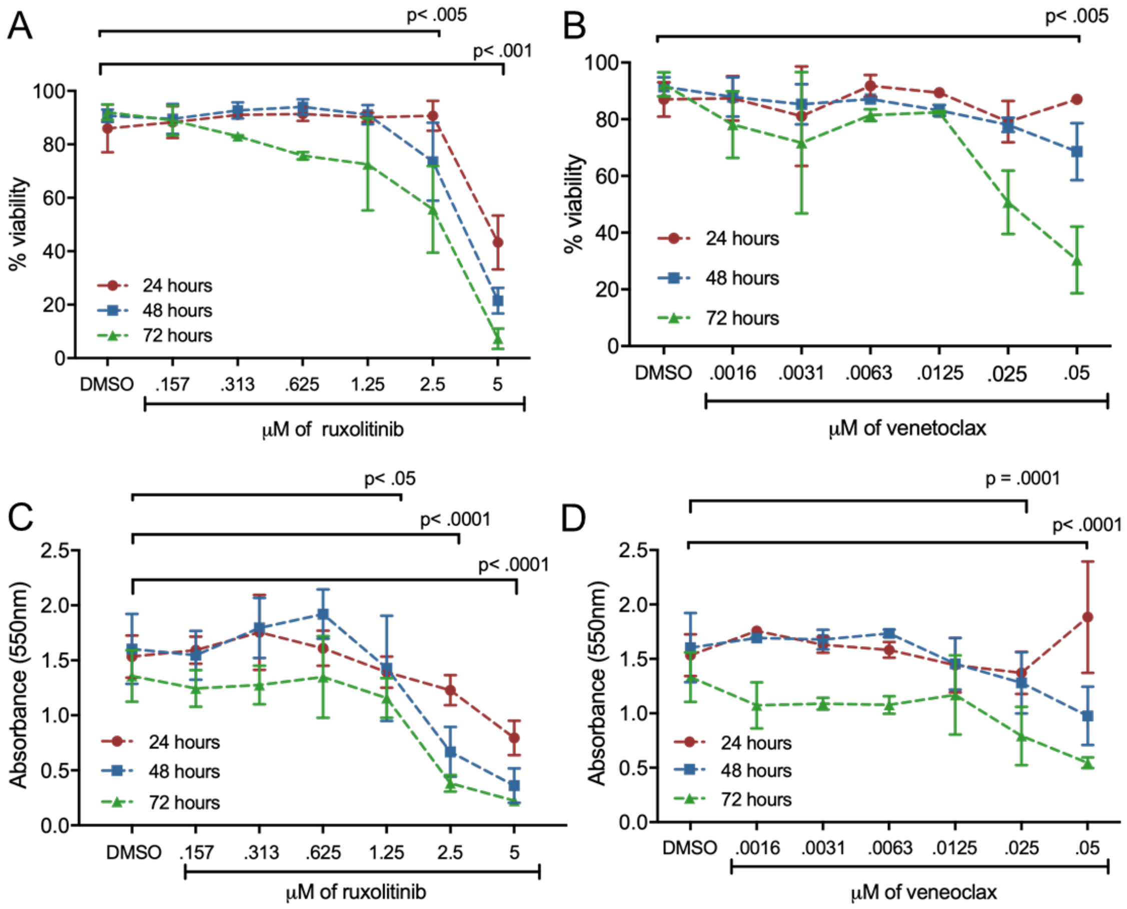


**Supplemental Figure 1**

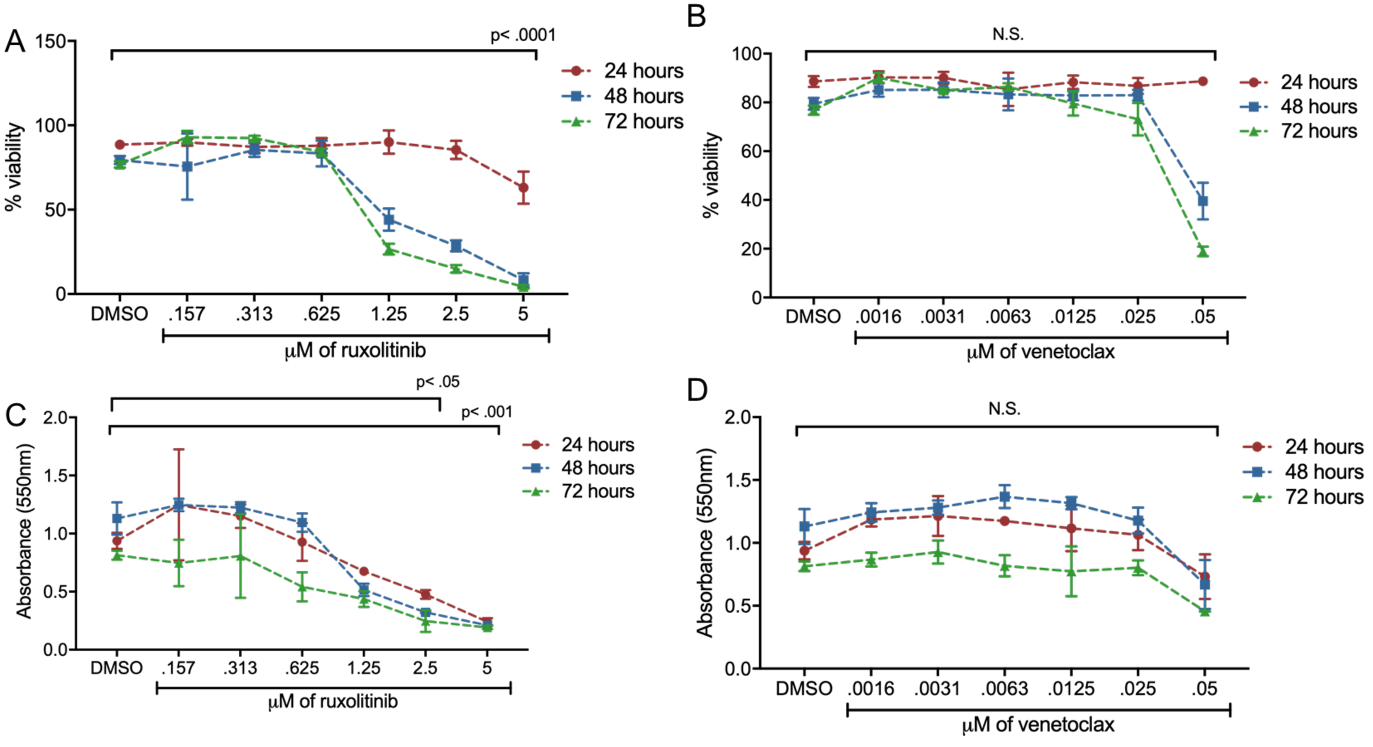


**Supplemental Figure 2**

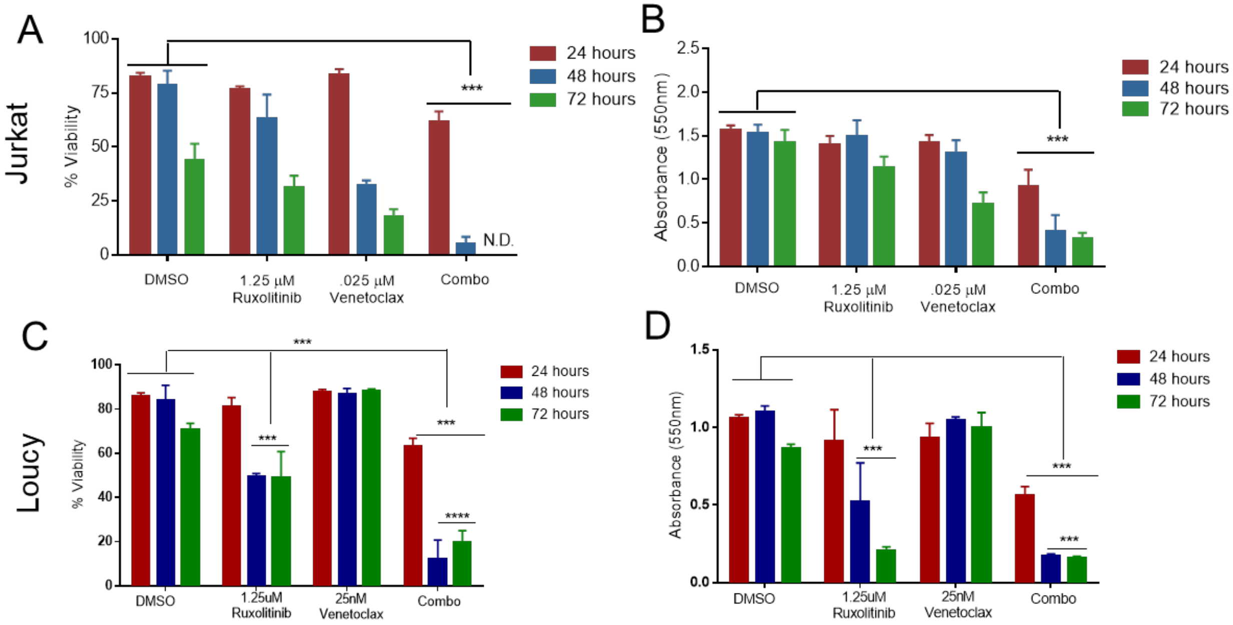


**Supplemental Figure 3**

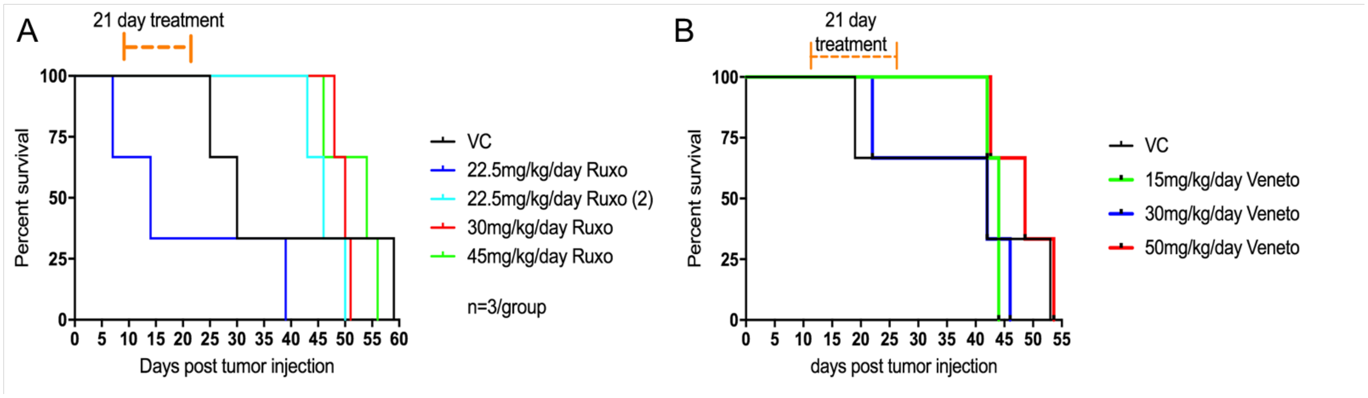


**Supplemental Figure 4**

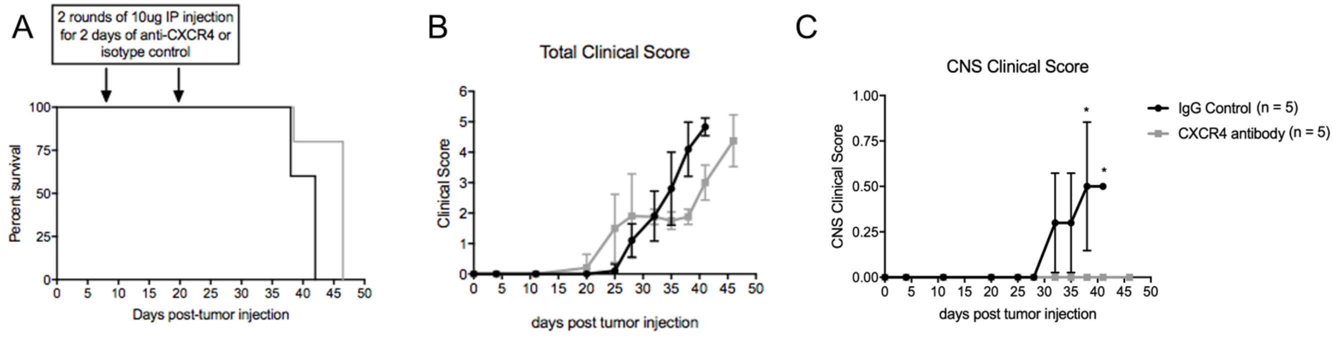


**Supplemental Figure 5**

**Supplementary Table 1. T-ALL clinical scoring criteria**

| | Score | | | | |
|---|---|---|---|---|---|
| **Criteria** | **0** | **0.5** | **1** | **1.5** | **2** |
| Activity | Normal | Mildly Decreased | Moderately Decreased | Stationary unless stimulated | Stationary even when stimulated; labored breath. |
| Weight loss | <10% | 10-15% | 15-20% | 20-25% | 25-30% |
| Hunch | Normal | Hunched when stationary; goes away when active | Small permanent hunch | Moderate permanent hunch | Severe hunch |
| Appearance | Normal | Slight ruffle | Moderate ruffle | Ruffle/ Greasy | Grizzled/Frizzy |
| Paralysis | Normal | Slow moving, less tension in tail upon lifting | partial, mouse starts biting/scratching hind legs | One leg completely paralyzed. Other has partial mobility | Both legs are completely paralyzed (hind-limbs dragging) |

Mice are euthanized when a total clinical score of 4 or higher is reached, or if mouse develops full paralysis (score of 2) or surpasses 25% weight-loss
